# Genotypic and Phenotypic Diversity of *Kluyveromyces marxianus* Isolates Obtained from the Elaboration Process of Two Traditional Mexican Alcoholic Beverages Derived from Agave: Pulque and Henequen (*Agave fourcroydes*) Mezcal

**DOI:** 10.1101/2023.07.04.547527

**Authors:** Patricia Lappe-Oliveras, Morena Avitia, Sara Darinka Sánchez-Robledo, Ana Karina Castillo-Plata, Lorena Pedraza, Guillermo Baquerizo, Sylvie Le Borgne

## Abstract

Seven *Kluyveromyces marxianus* isolates from the elaboration process of pulque and henequen mezcal were characterized. The isolates were identified based on the sequences of the D1/D2 domain of the 26S rRNA gene and the internal transcribed spacer (ITS-5.8S) region. Genetic differences were found between pulque and henequen mezcal isolates and within henequen mezcal isolates, as shown by different branching patterns in the ITS-5.8S phylogenetic tree and (GTG)5 microsatellite profiles, suggesting that substrate and process selective conditions may originate different *K. marxianus* populations. All the isolates fermented and assimilated inulin and lactose and some henequen isolates could also assimilate xylose and cellobiose. Henequen isolates were more thermotolerant than pulque isolates which in contrast presented more tolerance to the cell wall-disturbing agent calcofluor white (CFW) suggesting that they had a different cell wall structure. Additionally, depending on their origin, the isolates presented different maximum specific growth rates (µ_max_) patterns at different temperatures. Concerning tolerance to stress factors relevant for lignocellulosic hydrolysates fermentation, tolerance limits were lower at 42 than 30°C except for glucose and furfural. Pulque isolates were less tolerant to ethanol, NaCl, and Cd. Finally, all the isolates could produce ethanol by simultaneous saccharification and fermentation (SSF) of a corncob hydrolysate under laboratory conditions at 42°C.

## 1. Introduction

The nonconventional yeast *Kluyveromyces marxianus* has attractive characteristics for industrial applications as a high growth rate, thermotolerance, a broad spectrum of carbon sources utilization including xylose, lactose and inulin, and the ability to produce ethanol, aroma compounds, enzymes, vaccines, bioactive molecules, fructo-oligosaccharides, fatty acids, and single-cell proteins [1–5]. *K. marxianus* has also being considered for use as a probiotic [6]. Applications in wine production have also been suggested [7]. This yeast is a promising cell factory for biorefinery applications using lignocellulosic biomass hydrolysates and dairy industry lactose-rich effluents as feedstocks [8–10]. Due to its thermotolerance, *K. marxianus* has been considered of interest for the simultaneous saccharification and fermentation (SSF) of lignocellulosic biomass hydrolysates in which the fermentation must be carried out at high temperatures (>40ºC) close to the optimal temperature for enzymatic saccharification (50°C) [11–13].

*K. marxianus* has most frequently been isolated from dairy environments as raw milk [14], cheese as Pecorino di Farindola [15] and traditional fermented milk products from different parts of the world as kefir [16], among others. This association with dairy products is due its capability to metabolize lactose. This species has also been isolated from plants and fruits as, agave [17], rotting onions [18], overripened mango pulp [19] and agro-industrial residues in sugar mills [20], sugarcane bagasse hydrolysates [21], coffee wet processing wastewater [22], blue agave bagasse [23], and distillery effluents and molasses [24]. *K. marxianus* is also part of the microbiota involved in the fermentation of cereal-based African fermented beverages [25], French cider [26], Georgian wine [27], Brazilian cachaça [28], and Mexican agave-based spirits tequila and mezcal [29–31].

As emphasized by several authors, *K. marxianus* exhibits a substantial genetic and physiological diversity as illustrated by the variety of habitats from which it can be isolated [1,32,33]. This diversity has mainly been studied in strains isolated from dairy products [15,34–36]. However, it has been recently proposed that *K. marxianus* strains isolated from agave or associated with agave-based fermentations may represent a divergent clade compared to strains from dairy environments and other habitats [37].

The objectives of the present study were to genotypically and phenotypically characterize *K. marxianus* isolates obtained from the elaboration process of pulque and henequen mezcal, two agave-based alcoholic beverages, as well as to evaluate their tolerance to different stress conditions, and their ability to produce ethanol from a corncob hydrolysate by a SSF procedure.

Pulque is an ancient Mesoamerican non-distilled beverage (4-7% of ethanol) made from several agave species cultivated in the Central Mexican plateau [38]. The plant is not cooked, instead, the fresh plant sap (*aguamiel*) is extracted directly from the plant by scrapping the cavity made in the center of the agave stem. The obtained sap is fermented for 24 h to several days by adding a portion of previously fermented pulque (called seed or *semilla*) as inoculum, and directly consumed after the fermentation process has finished [39]. Mezcal is a spirit distillate made from the stems or cores (called *piñas*) of a variety of agave species, which are slowly cooked in large pit ovens, then crushed in stone mills to obtain a sweet juice that is fermented and finally distilled [40]. Henequen (*Agave fourcoydes*) is an agave species native to the Yucatan Peninsula [41].

## 2. Materials and Methods

### 2.1. Yeasts strains and growth conditions

Seven isolates of *K. marxianus* recovered from different stages of the production process of pulque and henequen mezcal were obtained from the Microbial Culture Collection (yeasts and molds) at the Laboratorio de Micromicetos (C006) of the Instituto de Biología of the Universidad Nacional Autónoma de México (Table 1). The artisanal pulque was produced in Santa Mónica (Hidalgo State, Mexico: 20°18′31.542″N, 98°13′8.076″W) [42], and henequen mezcal at the GeMBio Laboratory, Centro de Investigación Científica de Yucatán A.C. in Merida, Yucatan State, Mexico [43] from samples collected in Tixpéhual (Yucatán State, Mexico: 20 57 39.321 N, 89 26 21.671 W). Axenic yeast cultures were preserved in aqueous suspensions at 4°C and in YPD broth (20 g/L bacteriological peptone, 10 g/L yeast extract, 20 g/L of glucose) added with 25% (v/v) of glycerol at -80°C. Unless otherwise indicated, cultures were performed in YPD broth or in YPD agar and incubated at 30°C.

**Table 1.** Origin of the *K. marxianus* isolates. Circles at the left of each isolate key are colored according to their origin.

| Isolate | Origin |
| --- | --- |
| ● Kmx11 | Pulque seed or inoculum |
| ● Kmx14 | Base of a freshly cut henequen leaf |
| ● Kmx15 | Fermented (168 h) pulque |
| ● Kmx16 | Non-fermented cooked henequen juice |
| ● Kmx21 | Non-fermented cooked henequen juice |
| ● Kmx22 | Fermented (48 h) cooked henequen juice |
| ● Kmx24 | Cooked henequen stem |

### 2.2. DNA extraction, molecular identification, and phylogenetic analysis

Genomic DNA was extracted using the ZR Fungal/bacterial DNA Miniprep kit (Zymo Research Corp, Irvine, CA, USA) according to the manufacturer^’s^s instructions. The concentration and purity of the DNA was determined using the NanoDrop 2000 spectrophotometer (Thermo Scientific, Waltham, MA, USA) with absorbance readings at 230, 260 and 280 nm. DNA integrity was evaluated by electrophoresis in 0.8% (w/v) agarose gels stained with GelRed (Biotium, Fremont, CA, USA).

The D1/D2 domain of the 26S rRNA gene was amplified using the forward NL1 (5^’s^-GCA TAT CAA TAA GCG GAG GAA AAG-3’) and reverse NL4 (5’-GGT CCG TGT TTC AAG ACG G-3’) primers [44] in 25-µL PCR reaction volume containing 25 ng of DNA template, 1x standard Taq reaction buffer (New England Biolabs, Ipswich, MA, USA), 0.2 mM of each dNTP, 0.5 µM of each primer and 1.25 U Taq DNA polymerase (New England Biolabs). DNA amplification was performed in a MyCycler thermal cycler (BioRad, Hercules, CA, USA) under the following conditions: 5 min of initial denaturation at 95°C ; 30 cycles of 30 s at 94°C, 30 s at 55°C and 30 s at 72°C; one cycle of final extension of 7 min at 72°C.

The 5.8S-ITS region of the 26S rRNA gene was amplified using the forward ITS1 (5^’s^-TCC GTA GGT GAA CCT GCG G-3’) and reverse ITS4 (5’-GCA TAT CAA TAA GCG GAG GA-3’) primers [45] in 25-µL PCR reaction volume as described above. DNA amplification was performed under the following conditions: 5 min of initial denaturation at 95°C; 35 cycles of 1 min s at 94°C, 2 min at 55.5°C and 2 min at 72°C; one cycle of final extension of 10 min at 72°C.

All the amplicons were analyzed by electrophoresis in 1.2% (w/v) agarose gels stained with GelRed. After purification with the DNA Clean and Concentrator kit (Zymo Research), the D1/D2 and ITS-5.8S PCR amplification products were sent to Macrogen (Seoul, Korea) for sequencing. The D1/D2 sequences were edited with BioEdit 7.0.5 [46] and compared with the sequences in the GenBank database using the BLASTN online tool [47] for identification. For the ITS-5.8S phylogeny analysis, sequences were aligned using BioEdit and the phylogenetic reconstruction was performed with the neighbor-joining (NJ) algorithm in MEGA7 [48] using 10,000 bootstrap replicates.

### 2.3. Microsatellites analysis

Microsatellite PCR fingerprinting was carried out in 25-µL reaction volume containing 25 ng of DNA template, 1x standard Taq reaction buffer (New England Biolabs), 0.2 mM of each dNTP, 0.5 µM of the (GTG)5 microsatellite primer, 2.5 μg of bovine serum albumin (BSA) and 1.25 U Taq DNA polymerase (New England Biolabs) according to [49]. DNA amplification was performed in a MyCycler thermal cycler (Bio-Rad) under the following conditions: 5 min of initial denaturation at 94°C; 40 cycles of 30 s at 94°C, 45 s at 55°C and 1 min and 30s at 72°C; one cycle of final extension of 6 min at 72°C.

The amplification products were separated on 2% (w/v) agarose gels containing GelRed in 0.5X TBE buffer at 7.5 V/cm for 150 min. A 100 pb DNA ladder (New England Biolabs) was used as the molecular weight marker in these gels.

### 2.4. Morphological and phenotypical characterization

The colonial and ascospore morphology as well as carbon source utilization profiles were described and tested according to already described methodologies [50,51]. The morphological tests and most biochemical tests were performed at the Micromycetes Laboratory (C006) of the Institute of Biology of the Universidad Nacional Autónoma de México and some of the biochemical tests at the BCCM/MUCL culture collection (Université Catholique de Louvain, Belgium).

### 2.5. Thermotolerance and calcofluor white tolerance evaluation

Tolerance to temperature and calcofluor white (CFW) was tested by spot inoculation of liquid cultures previously grown at 30°C onto YPD agar as already described [33]. For this, the isolates were grown for 16 h at 30°C and 150 rpm in YPD broth and re-inoculated into fresh medium and grown to exponential phase at 30°C. The obtained cultures were adjusted to an optical density of 0.2 at 600 nm (OD_600_) in YPD broth, serially diluted into saline solution (NaCl 9% and spot inoculated (2.5 µL) onto the different test media and incubated at the temperatures listed below. The plates were sealed using parafilm before incubation and growth was observed after 24 and 48 h. For thermotolerance, the plates were incubated at 30, 37, 42, 45 and 48°C. For CFW tolerance, YPD agar was supplemented with 0.02, 0.05, 0.1, 0.15 or 0.2 mM of CFW and incubated at 30, 37, 42 and 45°C. The CFW (F3543, Sigma-Aldrich, Saint Louis, MO, USA) stock solution was prepared by dissolving CFW in 0.5% (w/v) KOH and 83% (v/v) glycerol according to [52]. The YPD agar was supplemented with this stock solution to obtain the CFW concentrations listed above.

### 2.6. Growth kinetics at different temperatures

Inocula were prepared in YPD broth from one colony inoculated in 5 mL of medium and grown at 30°C and 200 rpm for 16 h. The obtained cultures were used to inoculate 125 mL Erlenmeyer flasks containing 25 mL of YPD broth at an initial OD600 of 0.25. The flasks were then incubated at different temperatures (30, 37, 42, 45 and 48°C) for 24 h at 200 rpm. Cell growth was followed by measuring the OD600 in a Bio-Photometer Plus (Eppendorf, Hamburg, Germany).

The modified Gompertz model (Eq. (1)) was used to evaluate the effect of temperature on the shape of each isolate growth curve and determine the maximum growth rate (µ_max_) and the lag phase duration [53]. Model parameters were determined by fitting the growth curves to the Gompertz equation using the Levenberg-Marquardt nonlinear least squares method programmed in MATLAB^®^.

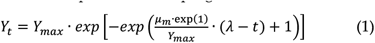

Where Y_t_ is the OD_600_ value at a time t, Y_max_ the maximum OD_600_ value, µ_m_ the maximum growth (OD_600_·h ^-1^), and λ the lag phase time (h). In addition, μ_max_ (h^-1^) was calculated with Eq. (2). The accuracy of the model was assessed through the coefficient of determination (R^2^), according to Eq. (3).

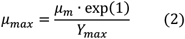

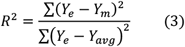

### 2.8. Tolerance to stress conditions relevant for lignocellulosic hydrolysates fermentation

Tolerance to stressful conditions was assessed by spot inoculation (as described in section 2.5) of liquid cultures inoculated onto YPD agar plates supplemented with the different test compounds and observing growth after 24-48 h [54]. *T*he plates were incubated at 30 and 42°C. The isolates were evaluated for tolerance (i) to glucose (2-50%, w/v); ii) to ethanol (2.5-10%, v/v); iii) to NaCl and KCl (25-75 g/L and 25-100 g/L, respectively); iv) to acetic acid (1.5-4.5 g/L), furfural (1-2.5 g/L) and coniferyl aldehyde (0.25-4 mM); and v) to metals (ZnCl_2_, 0.625–10 mM;, CuCl_2_, 0.1–2 mM; CdCl_2_, 0.25–1 mM; MnCl_2_, 2-6 mM).

### 2.9. SSF tests

The lignocellulosic substrate tested for bioethanol production was a pretreated corncob residue. The pretreatment consisted of a thermochemical treatment at a moderate temperature with diluted sulfuric acid to hydrolyze hemicellulose and the solid fraction primarily constituted by cellulose was used as substrate as described in [55]. SSF tests were conducted in duplicate in 125 mL Erlenmeyer flasks sealed with rubber stoppers. Each flask contained 8% (w/v) of pretreated corncob solids in a total volume of 62.5 mL of fermentation medium (5 g/L of yeast extract, 2 g/L of NH_4_Cl, 1 g/L of KH_2_PO_4_ and 0.3 g/L of MgSO_4_ 7H_2_O) without glucose and supplemented with 0.1 M of sodium citrate buffer at pH 5.5. The pH of the medium was finally adjusted to 5.5 with NaOH 2N after adding all the components. The pretreated corncob*s* and the medium were sterilized in autoclave prior to SSF. The *K. marxianus* isolates previously grown up to early *stationary* phase in YPD broth w*ere* inoculated at an initial OD_600_ of 0.5. 9% (w/w) of the commercial cellulase cocktail CelliCtec 2 (Novozymes Latin America, Araucária, Paraná, Brazil) with respect to the cellulose content were then added and the flasks were incubated at 42°C with an orbital agitation of 150 rpm for 72 h. Glucose, xylose, acetic acid, glycerol, and ethanol were determined by high-performance liquid chromatography (HPLC) as described in [56].

### 2.10. Nucleotide sequences accession numbers

The sequences obtained in this study were deposited in the GenBank database under the accession numbers shown in Table 2.

**Table 2.**
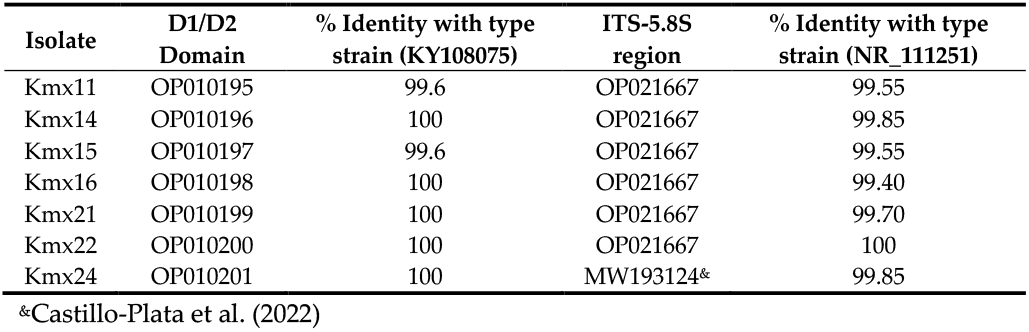
GenBank accession numbers and percent of nucleotide similarity in the D1/D2 domains and ITS regions between the isolates and type strain CBS 712.

## 3. Results

### 3.1. Molecular identification and phylogenetic analysis

The identity of the *K. marxianus* isolates was obtained by sequence analysis of the D1/D2 domain of the 26S rRNA gene. All sequences had a similarity value of 99.6 to 100% compared to the type strain sequence (*K. marxianus* CBS 712) (Table 2). Sequencing of the ITS-5.8S region confirmed the *K. marxianus* species as-signation of all the isolates with above 99% identity with *K. marxianus* CBS 712 (Table 2).

A phylogenetic tree was constructed with all the ITS-5.8S sequences of the showing that they grouped according to their origin (see Table 1 for the origin and color code of the different isolates) except Kmx14 from the base of a henequen leaf and Kmx24 from a cooked agave core which grouped together despite their different origin (Figure 1).

**Figure 1.**
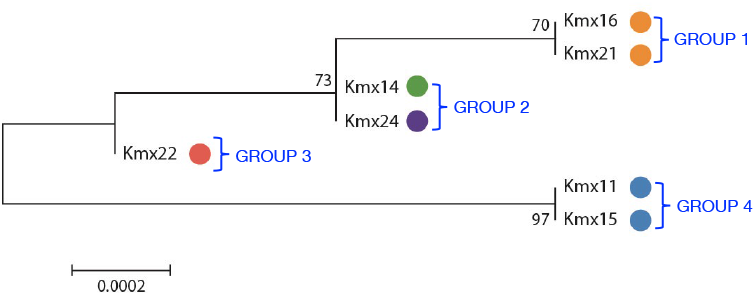
Phylogenetic tree of the *K. marxianus* isolates based on their ITS-5.8S sequences. The tree was constructed using 690 bp sequences. Colored circles indicate the origin of the isolates as described in Table 1. Numbers on nodes indicate bootstrap values. Branch lengths are proportional to the number of nucleotide substitutions and are measured using the bar scale (0.0002). Sequence accession numbers are shown in Table 2.

### 3.2. Microsatellites

To assess the genetic diversity among the *K. marxianus* isolates, microsatellite PCR fingerprinting with the (GTG)_5_ primer was performed. The observed patterns were simple and consisted of 3 to 5 bands with sizes between 500 and 2,000 bp (Figure 2). As in theITS-5.8S phylogenetic tree, the isolates clustered according to their origin except for Kmx14 and Kmx24. The pulque isolates Kmx11 and Kmx15 produced the same (GTG)_5_ banding pattern (pattern I in Figure 2 and Group 4 in Figure 1), while the henequen mezcal isolates clustered into three different (GTG)_5_ banding patterns: pattern II in Figure 2 (Group 2 in Figure 1) for Kmx14 and Kmx24 isolated from a henequen leaf base and from a cooked agave stem, respectively; pattern III in Figure 2 (Group 1 in Figure 1) for Kmx16 and Kmx21, both from non-fermented juice extracted from cooked henequen stems; and pattern IV (Group 3 in Figure 1) for Kmx22 from fermented (48 h) henequen juice.

**Figure 2.**
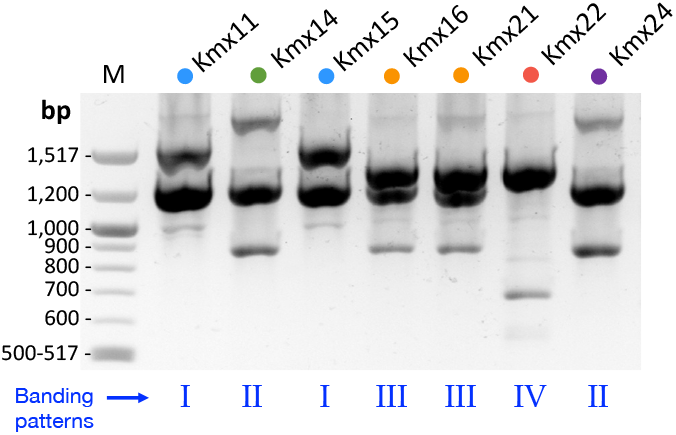
PCR-fingerprinting of the *K. marxianus* isolates obtained with the (GTG)^5^ microsatellite primer.

### 3.3. Morphology and carbon sources utilization

All strains formed cream-colored colonies in YPD agar except Kmx16 which presented a pinkish coloration. Concerning ascospore shape, all isolates formed reniform ascospores except Kmx22 which produced round ones. Table 3 presents the results of the carbon sources fermentation and assimilation tests of compounds for biotechnological applications. Table S1 shows additional carbon and nitrogen assimilation tests. All the isolates were able to grow at 37°C, ferment glucose, assimilate raffinose and did not assimilate maltose, which are the key characteristics of *K. marxianus* according to [57]. All the isolates were clearly positive for lactose, inulin, and xylitol assimilation and could ferment galactose, lactose, and inulin but not xylose and cellobiose. Positive, delayed, or weak assimilation profiles were recorded for xylose and cellobiose while negative, delayed, or weak profiles were observed for sugar alcohols other than xylitol, gluconolactone and citrate. Finally, a positive or weak succinate assimilation pattern was found instead of the positive or delayed profiles reported for this species. All the isolates could assimilate lactate, but Kmx24 gave a weak instead of a positive response. A positive ethanol assimilation was also observed for all the isolates, as expected for *K. marxianus*.

**Table 3.** Carbon sources fermentation and assimilation patterns of the *K. marxianus* isolates. The two last columns show the results reported for *K. marxianus* in the literature for comparison.

| Carbon source | Isolate |  |  |  |  |  |  | [58] | [59] |
| --- | --- | --- | --- | --- | --- | --- | --- | --- | --- |
|  | K<br>m<br>x1<br>1 | Kmx<br>14 | Kmx<br>15 | Kmx<br>16 | Kmx<br>21 | Kmx<br>22 | Kmx<br>24 |  |  |
| Fermentation |  |  |  |  |  |  |  |  |  |
| Glucose | + | + | + | + | + | + | + | + | + |
| Fructose | + | + | + | + | + | + | + | n.d. | n.d. |
| Galactose | + | w | + | + | + | + | + | d | +, d |
| Xylose | - | - | - | - | - | - | - | n.d. | - |
| Sucrose | + | + | + | + | + | + | + | + | + |
| Maltose | - | - | - | - | - | - | - | - | +, - |
| Lactose | + | + | + | + | + | + | + | v | +, - |
| Cellobiose | - | - | - | - | - | - | - | n.d. | d, - |
| Raffinose | + | + | + | + | + | + | + | + | +, - |
| Inulin | + | + | + | + | + | + | + | d | +, - |
| Assimilation |  |  |  |  |  |  |  |  |  |
| Glucose | + | + | + | + | + | + | + | + | + |
| Raffinose | + | + | + | + | + | + | + | + | + |
| Maltose | - | - | - | - | - | - | - | +, - | - |
| Rhamnose | - | - | - | - | - | - | - | - | - |
| Sucrose | + | + | + | + | + | + | + | + | + |
| Xylose | d | + | w | + | + | + | + | v | +, - |
| Ribose | - | w | - | - | - | w | d | v | +, - |
| Lactose | + | + | + | + | + | + | + | v | +, - |
| Cellobiose | + | + | + | + | + | d | + | v | +, - |
| Trehalose | - | - | - | - | - | - | - | - | +, - |
| Inulin | + | + | + | + | + | + | + | v | +, - |
| Glycerol | - | w | w | - | - | w | w | v | +, d |
| Ribitol | d | w | d | - | D | w | d | v | +, - |
| Xylitol | + | + | + | + | + | + | + | v | +, - |
| Mannitol | - | w | w | - | W | W | w | v | +, - |
| Sorbitol | d | + | + | w | D | W | + | v | +, - |
| Gluconolactone | - | - | - | - | - | - | - | v | +, - |
| Citrate | - | - | - | w | - | - | - | v | +, - |
| Succinate | w | w | w | + | + | + | w | + | +, d |
| DL-Lactate | + | + | + | + | + | + | w | + | + |
| Ethanol | + | + | + | + | + | + | + | + | +, d |

+, positive; -, negative; d, delayed: positive response > 14 days; w, weak positive response; n.d, not determined. \*, positive response after a week

### 3.4. Thermotolerance and growth profiles at different temperatures

Growth of all isolates was tested at 30, 37, 42, 45, and 48 °C in YPD agar plates (Figure 3). All the isolates failed to grow at 48 °C under these conditions. Kmx14, Kmx16, Kmx21, Kmx22 and Kmx24 could grow up to 45°C, but Kmx22 presented less growth at this temperature. Kmx11 and Kmx15, isolated from pulque grew well up to 42°C.

**Figure 3.**
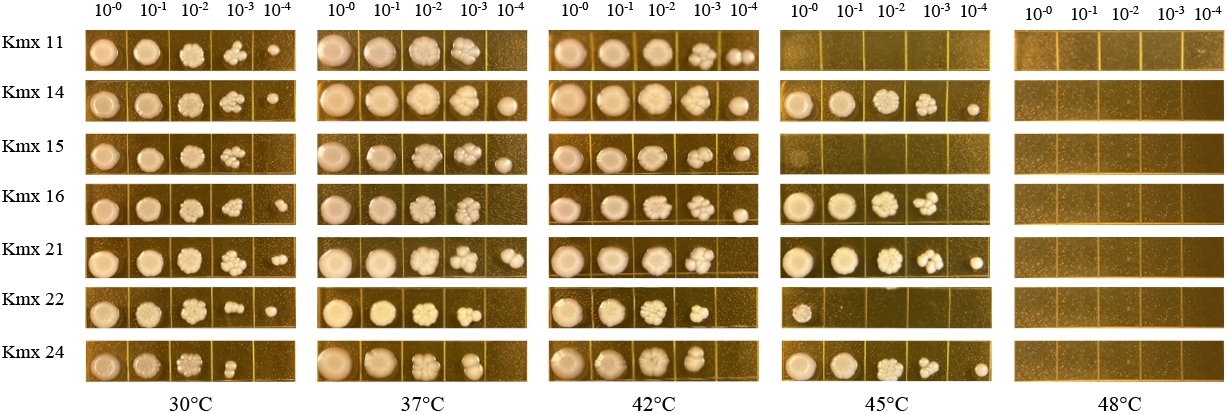
Growth of the *K. marxianus* isolates at different temperatures after 48 h of incubation in YPD agar. The dilutions are indicated on the top of the pictures.

The experimental growth curves of the isolates cultured in YPD broth at different temperatures were fitted to the modified Gompertz model, using growth data up to 42 and 45 °C for the pulque and henequen isolates, respectively. The calibrated parameters for each fermentation are detailed in Table S2, where the relative standard deviation for both µ_max_ and lag phase duration (λ) showed average values of 9.9% and 5%, respectively, demonstrating that the kinetic parameters were accurately determined. Figure 4 displays the µ_max_ of the isolates as a function of temperature (*i*.*e*. the thermal growth curves). Lower µmax values were observed for the pulque isolates (Figure 4A) compared to the henequen isolates (Figure 4B, 4C and 4D). The optimum growth temperatures of pulque isolates (37-42 °C) were also lower than that of henequen isolates (42 °C). Isolate Kmx22 from henequen showed a different behavior with an almost flat pattern over the range of tested temperatures and intermediate µmax values with respect to the pulque and henequen isolates (Figure 4D). The shape of the thermal curves was similar for isolates belonging to the same ITS-5.8S group/microsatellite pattern.

**Figure 4.**
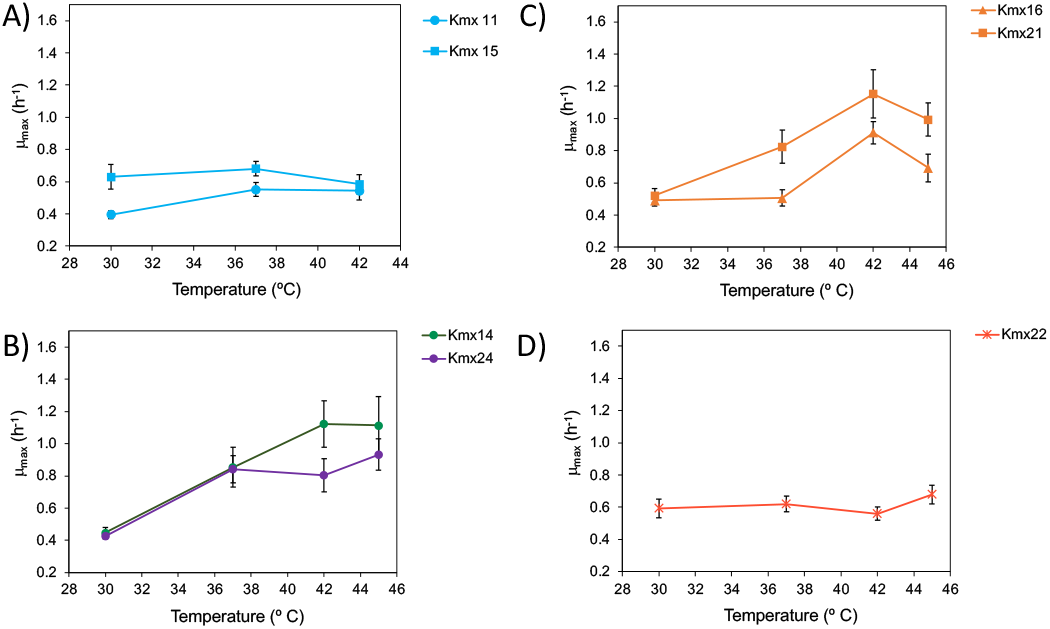
Plots of maximum growth rate (µ_max_) of the isolates versus temperature. A) Pulque isolates; B, C and D) Henequen isolates grouped according to their ITS-5.8S group/microsatellite pattern.

### 3.5. Tolerance to CFW at different temperatures

Tolerance to the cell wall perturbing agent CFW was tested to evaluate possible differences in the cell wall structure among the isolates and at different temperatures. CFW interferes with the cell wall assembly by binding to chitin [52] and has been used to measure the chitin content of yeast cell walls [60]. Cells with a high chitin content bind more CFW and have a low tolerance to this dye and a high staining index. The growth results are shown in Figure 5. Most of the isolates were sensitive to CFW and, in general, good growth was observed with only 0.05 mM of this compound at 30 °C, except for Kmx11 and Kmx15 which were clearly more tolerant and could grow at up to 0.2 and 0.5 mM CFW at this temperature. Kmx22 was the least tolerant isolate and presented a limited growth at the lowest CFW concentration tested (0.02 mM) at 30°C. In all cases, the tolerance to CFW decreased at higher incubation temperatures and Kmx11 and Kmx15 were the most tolerant isolates at 42 °C.

**Figure 5.**
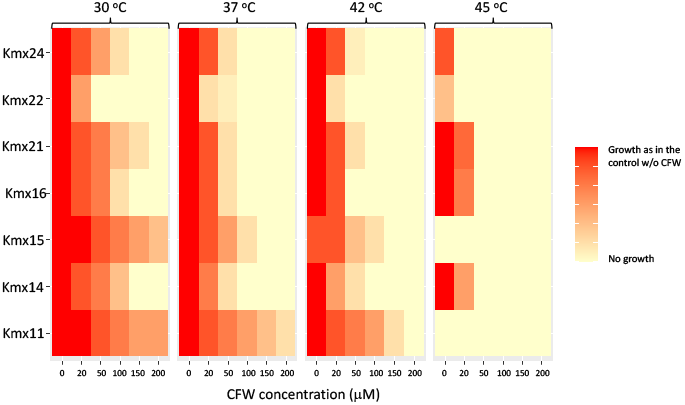
Tolerance to CFW at different temperatures.

### 3.6. Tolerance to stress conditions relevant for bioethanol production

Table 4 shows the tolerance limits of the *K. marxianus* isolates for glucose, ethanol, salts, fermentation inhibitors and metals, both at 30 and 42°C. In general, the isolates were tolerant to high concentrations of glucose (30%) at both temperatures except one isolate from pulque (Kmx15) which was less tolerant (15%) at 42 °C. All the isolates were less tolerant to ethanol (2.5-7.5%) at 42 than at 30 °C (7.5-10%) with Kmx11 and Kmx15 from pulque being the least tolerant. These two isolates did not grow in the presence of NaCl at 42 °C, while Kmx22 from fermented cooked henequen must was the most tolerant isolate (50g/L) at 30 °C. Concerning KCl, again the two isolates from pulque presented the lowest tolerance limit (50g/L) to this salt at 42 °C.

**Table 4.** Tolerance limits of the *K. marxianus* isolates to different concentrations of GLU, glucose; EtOH, ethanol; AA, acetic acid; FUR, furfural; CA, coniferyl aldehyde, at 30 and 42 °C. The concentrations tested are described in section 2.8.

| Isolate | GLU<br>(% w/v) |  | EtOH<br>(% v/v) |  | NaCl<br>(g/L) |  | KCl<br>(g/L) |  | AA<br>(g/L) |  | FUR<br>(g/L) |  | CA<br>(mM) |  | Zn<br>(mM) |  | Cd<br>(mM) |  | Cu<br>(mM) |  | Mn<br>(mM) |  |
| --- | --- | --- | --- | --- | --- | --- | --- | --- | --- | --- | --- | --- | --- | --- | --- | --- | --- | --- | --- | --- | --- | --- |
|  | 30°C | 42°C | 30°C | 42°C | 30°C | 42°C | 30°C | 42°C | 30°C | 42°C | 30°C | 42°C | 30°C | 42°C | 30°C | 42°C | 30°C | 42°C | 30°C | 42°C | 30°C | 42°C |
| • Kmx11 | 30 | 30 | 7.5 | 2.5 | 25 | 0 | 75 | 50 | 3 | 0 | 2 | 2 | 2 | 1 | 5 | 2.5 | 0.25 | 0 | 4 | 2 | 2 | 2 |
| • Kmx14 | 30 | 30 | 10 | 7.5 | 37.5 | 37.5 | 75 | 75 | 3 | 1.5 | 1.5 | 2 | 2 | 2 | 5 | 2.5 | 0.5 | 0.25 | 4 | 4 | 2 | 2 |
| • Kmx15 | 30 | 15 | 5 | 2.5 | 25 | 0 | 75 | 50 | 3 | 1.5 | 2 | 2 | 2 | 1 | 5 | 1.25 | 0.25 | 0 | 4 | 2 | 2 | 0 |
| • Kmx16 | 30 | 30 | 10 | 5 | 37.5 | 37.5 | 75 | 75 | 3 | 1.5 | 1.5 | 2 | 2 | 2 | 5 | 2.5 | 0.5 | 0.25 | 4 | 4 | 2 | 2 |
| • Kmx21 | 30 | 30 | 10 | 5 | 37.5 | 37.5 | 75 | 75 | 3 | 1.5 | 1.5 | 2 | 2 | 2 | 5 | 2.5 | 0.5 | 0.25 | 4 | 4 | 2 | 2 |
| • Kmx22 | 30 | 30 | 7.5 | 5 | 50 | 37.5 | 75 | 75 | 3 | 1.5 | 1.5 | 1.5 | 2 | 2 | 2.5 | 2.5 | 0 | 0 | 4 | 4 | 4 | 2 |
| • Kmx24 | 30 | 30 | 7.5 | 5 | 37.5 | 37.5 | 75 | 75 | 3 | 1.5 | 1.5 | 2 | 2 | 2 | 5 | 2.5 | 0.5 | 0.25 | 4 | 4 | 2 | 2 |

All the isolates tolerated 3 g/L of acetic acid at 30 °C and were all less tolerant to this acid (1.5g/L) at 42 °C. In this case, Kmx11 from pulque did not grow in the presence of acetic acid at 42 °C. Contrary to acetic acid, the isolates were slightly more tolerant to furfural at 42 (1.5-2.0g/L) than at 30 °C, except the pulque isolates that equally tolerant to furfural at 30 and 42 °C. In the case of coniferyl aldehyde, contrary to furfural, the isolates from henequen were equally tolerant (2 mM) at both temperatures while pulque isolates presented the lowest tolerance (1 mM) at 42 °C. Concerning Zn tolerance, Kmx22 from henequen and Kmx15 from pulque were the less tolerant at 30 (2.5 mM) and 45 °C (1.25 mM), respectively. For Cd, all henequen isolates, were less tolerant at 42 °C (0.25 mM) than at 30 °C (0.5 mM) except Kmx22 which did not tolerate any of the Cd concentrations tested at both temperatures. Pulque isolates Kmx11 and Kmx15 were less tolerant at 30 (0.25 mM) than at 42 °C. Concerning Cu all the isolates showed the same tolerance (4 mM) at both temperatures tested, except for the pulque isolates that presented a lower tolerance at 42 °C (2 mM).

### 3.8. SSF of a corncob hydrolysate

The thermotolerance observed for the seven isolates allowed to perform SSF tests at 42 °C, the usual temperature of SSF with *K. marxianus* [11]. The unwashed solid from thermochemically pretreated corncob was used as substrate. The solid contained 63% of cellulose on a dry weight basis corresponding to a potential concentration of glucose of 55.8 g/L.

The initial concentration of acetic acid in the culture medium was 3.1 g/L. The ethanol, acetic acid and glycerol production and glucose consumption were determined at the end of the SSF trials (72 h) (Table 5). All the *K. marxianus* isolates were able to convert glucose into ethanol, with isolate Kmx11 being the most productive one (15.8 g ethanol/L) followed by Kmx21, Kmx24, Kmx22, Kmx16, Kmx14, and Kmx15. The increase in ethanol production was accompanied by a decrease in glucose concentration, which was almost zero at the end of the trial except for isolate Kmx15 in which the glucose concentration remained high (15 g/L). In the case of the most and less productive isolates (Kmx11 and Kmx15, both from pulque), ethanol yields were 56.4 and 43.3%, respectively while for Kmx14, Kmx16, Kmx21, Kmx22, and Kmx24 the calculated ethanol yields were 45.9 to 49.4% (Table 5).

**Table 5.** Ethanol yields and glucose (GLU), acetic acid (AA), glycerol (GLY), and ethanol (ETOH) concentrations obtained in the SSF of a corncob hydrolysate with the *K. marxianus* isolates.

| Isolate | GLU at 72 h<br>(g/L) | AA at 72<br>h (g/L) | GLY at 72<br>h (g/L) | EtOH at 72<br>h (g/L) | $Y_{P/S}^{(a)}$ | EtOH yield<br><sup>(b)</sup> |
| --- | --- | --- | --- | --- | --- | --- |
| Kmx11 | 0.8 | 3.5 | 0.8 | 15.8 | 0.3 | 56.4% |
| Kmx14 | 3.7 | 3.1 | 0.6 | 12.5 | 0.2 | 47.0% |
| Kmx15 | 15.0 | 3.1 | 0.2 | 9.0 | 0.2 | 43.3% |
| Kmx16 | 0.3 | 3.4 | 0.7 | 13.0 | 0.2 | 45.9% |
| Kmx21 | 0.4 | 3.3 | 0.5 | 13.9 | 0.3 | 49.3% |
| Kmx22 | 0.3 | 3.4 | 0.9 | 13.8 | 0.2 | 48.8% |
| Kmx24 | 0.6 | 3.6 | 0.5 | 13.9 | 0.3 | 49.4% |

(a) Yield product/substrate, g ethanol/g (Potential GLU in pretreated corccob –GLU at 72 h).
(b) As percentage of the maximum theoretical ethanol yield: 0.51 g ethanol/g glucose.

## 4. Discussion

Here, the genotypic and phenotypic characterization of two *K. marxianus* isolates from the elaboration process of pulque (Kmx11 and Kmx15) and five *K. marxianus* isolates from the elaboration process of henequen mezcal (Kmx14, Kmx16, Kmx21, Kmx22 and Kmx24) is reported (see Table 1 for a description of the precise origin of the isolates). It has been suggested that strains with more than 1% nucleotide substitutions in their D1/D2 domain sequence are most likely to belong to different yeast species, while strains with 1% or less substitutions are conspecific or sister species [61]. Therefore, according to the base pair sequence analysis of the D1/D2 domain of the 26S rRNA gene, all the isolates belonged to the *K. marxianus* species (Table 2). The identity of all isolates was further confirmed by sequencing the ITS-5.8S region (Table 2).

The phylogenetic tree constructed with the ITS-5.8S sequences separated the isolates into four groups, most of them in accordance with their origin (Figure 2), except Kmx14 isolated from the base of a henequen leaf and Kmx24 from cooked agave stem which clustered in the same group in the phylogenetic tree (Figure 2, group 2). As agave stems are obtained after cutting the leaves and comprise both the stem of the plant and leaves bases, it is somehow not surprising that isolates Kmx14 and Kmx24 grouped together, although Kmx24 was isolated from a cooked agave stem and not from a fresh plant. The ITS-5.8S region was therefore a useful marker for the differentiation of agave isolates within the *K. marxianus* species and showed that a significant intraspecific genetic diversity was present among these isolates. By examining the-ITS-5.8S sequences of eleven *K. marxianus* strains from different origins, [62] observed that four different ITS-5.8S sequence haplotypes were present among these strains, pointing out the high intraspecific diversity within this species. On the other hand, polymorphism in the ITS-5.8S sequence was discriminative enough to distinguish terroir *Saccharomyces cerevisiae* wine yeasts with specific fermentative properties [63].

In addition to the ITS-5.8S analysis, the genetic diversity of the isolates was also assessed by microsatellites PCR fingerprinting. Microsatellites are short DNA motifs repeated in tandem present in eukaryotic genomes. Although originally designed to study genetic variations in humans due to their high degree of variability, microsatellites have also become a powerful tool to study intraspecific diversity in yeasts, enabling for example to discriminate between *S. cerevisiae* strains from wine and beer [64] or from artisanal versus industrial bread-making processes [65]. Interestingly, microsatellites analysis of the *K. marxianus* isolates from henequen and pulque produced four different patterns (Figure 2) that corresponded to the four groups already detected in the ITS-5.8S phylogenetic tree (Figure 2) suggesting a link between the genotype and the substrate of origin. The pulque isolates were discriminated from the henequen isolates which were in turn separated into three distinct populations according to their substrate of origin.

The colony and spore morphologies were homogenous among the isolates except for Kmx16 and Kmx22 respectively. Kmx16, from non-fermented cooked henequen juice, presented a reddish color on YPD agar. This color has been attributed to the production of the siderophore pulcherrimin [58]. Kmx22, from fermented cooked henequen must, formed round instead of reniform ascospores as the rest of the isolates. Both reniform and round ascospores have been described in *K. marxianus* [58,59]. Ascospore shape has been used as an important character in ascomycetous yeast taxonomy, but its functional role has not yet been studied in detail. One study with the budding yeast *Dipodascus albidus* suggests that spore shape may aid the dispersal and survival of yeasts, being favored reniform over round spores for an efficient release [66]. Here, the Kmx22 isolate from fermented henequen juice was the only isolate with round ascospores and, interestingly, it was located on a separate branch in the ITS-5.8S phylogenetic tree and produced a unique microsatellite pattern (Figures 1 and 2). More studies are needed to understand the possible biological meaning of these results.

Concerning carbon sources utilization, the biochemical tests consisted mainly in testing carbon sources relevant for possible industrial yeast processes. The fermentation tests results were almost homogenous among the isolates except for Kmx14 which presented a weak response for galactose. All the isolates were able to ferment lactose and inulin under the conditions described in [51] for yeasts systematic studies. Inulin and lactose fermentation are considered variable traits in *K. marxianus* (Table 3). Inulin is a type of fructose polymer (fructan) that serves as a storage carbohydrate in agave plants [67]. It is therefore not surprising that isolates from agave could use inulin. The ability to use inulin is due to the presence of extracellular inulinase enzymes that broke down inulin into fructose, an easily assimilable and fermentable sugar.

In the Kmx16 and Kmx24 isolates, the positive response for lactose fermentation was observed after a week, which could indicate that these two isolates were slow in fermenting lactose. Important differences in the kinetics of growth and ethanol production from lactose have been reported in *K. marxianus* strains from dairy or unknown environments, soil, fermented corn dough (pozol), and rooting agave leaf from South Africa. In particular, the agave strain CBS 745 presented the lowest biomass and ethanol yields on lactose [33]. Recent studies based on genomic analyzes indicate that the lactose fermentative metabolism of *K. marxianus* is related to the presence and expression of functional alleles of the lactose permease gene (LAC12) in dairy strains [37,68]. Detailed physiological and genomic studies are needed to characterize the lactose fermentative metabolism of the agave related isolates reported here.

Regarding carbon sources assimilation, the isolates showed a more phenotypic diversity (Table 3). All isolates could assimilate lactose, inulin and cellobiose while xylose was only efficiently assimilated by henequen derived isolates. This may not be surprising as agave fresh sap used for pulque elaboration mainly contains fructose, glucose, fructooligosaccharides and sucrose [69,70] while noticeable amounts of xylose are found in cooked agave juices used for mezcal elaboration [71]. The isolates related with henequen plant and mezcal elaboration may have adapted to a xylose containing environment. As mentioned above, inulin assimilation by all the isolates is not surprising as this sugar is present in agave plants. Cellobiose can be assimilated by some *K. marxianus* strains that possess a specific cellobiose transporter, or a dual lactose transporter, depending on the genetic background of the strains, and a cellobiase enzyme that hydrolyze cellobiose to glucose [72]. As xylose is a major component of hemicellulose, the ability to use this sugar as carbon source is relevant for lignocellulosic biomass utilization. Assimilation of cellobiose, a disaccharide produced during the partial hydrolysis of cellulose, is also relevant for the integral use of lignocellulosic biomass.

Concerning polyols assimilation, all the isolates could efficiently assimilate xylitol and some isolates sorbitol, ribitol, mannitol and glycerol under the conditions described in [51]. Polyols transport and metabolism in yeasts has been poorly studied although the ability of yeasts to assimilate polyols is part of the physiological tests for yeasts phenotypic characterization. According to the analysis performed in [73], most yeasts can assimilate at least one polyol, 10% of the described species can assimilate four polyols (arabitol, ribitol, sorbitol and xylitol) and ascomycetous yeasts preferably assimilate glycerol, followed by sorbitol and mannitol. As *K. marxianus*, most *Kluyveromyces* species give a variable response to xylitol [51,61]. This polyol is a low calory sweetener with a growing demand in the food sector. Both wild type and engineered *K. marxianus* strains have been used for xylitol production from lignocellulosic biomass and the main challenges to improve xylitol production in K. marxianus are related to xylose uptake and NADP supply [10]. However, it has also been reported that xylitol consumption or reassimilation by xylitol-producing yeasts also tends to reduce xylitol yields [74], so another limiting factor in the final xylitol yield and productivity may also be the reassimilation of the produced xylitol by *K. marxianus*. Finally, regarding carboxylic acids assimilation, as expected, all the isolates were clearly positive for DL-lactate except Kmx24 which showed a weak response. Six of the isolates were negative for citrate assimilation, with a weak response obtained for Kmx16. Three and four of the seven isolates gave a positive and weak response for succinate, respectively. The results obtained for these two Krebs substrates are consistent with the positive and delayed responses described in [58,59] confirming the Krebs-positive status of these *K. marxianus* isolates [75].

The ability of *K. marxianus* to grow at high temperatures is one of the remarkable characteristics of this species which is not present in the other *Kluyveromyces* species described so far. All the *K. marxianus* isolates reported here were able to grow at up to 42°C on YPD agar however, only the isolates from henequen grew at up to 45°C and none of the isolates could grow at 48°C (Figures 3). These temperature limits had already been reported for *K. marxianus* [33,36]. According to studies performed in *S. cerevisiae*, the physiological base of yeasts thermotolerance is complex and influenced by multiple genes [76]. Differences in thermotolerance limits have been frequently observed in industrial strains of *S. cerevisiae* [77] and linked to the presence of superior alleles in specific genomic *loci* in more thermotolerant strains [78]. No conserved thermo-tolerance mechanism has been found in thermotolerant yeasts [79]. It has been reported that an evolutionary young gene with unknown function was required for competitive growth of *K. marxianus* at high temperature [80]. However, this gene did not confer thermotolerance to *Kluyveromyces lactis* indicating that thermo-tolerance might be linked to the *de novo* emergence of species-specific genes.

The isolates exhibited different growth patterns in terms of optimum growth temperature, maximum growth rate, and thermal growth curve shape (Figure 4) and, interestingly, these differences corresponded to the groups and patterns observed in the ITS-5.8S and microsatellite patterns (Figures 1 and 2) indicating that the thermal adaptation of these yeasts was different according to their origin. For example, the isolates from pulque had a lower thermotolerance limit compared to all the henequen isolates and were located on a separate branch in the ITS-5.8 phylogenetic tree (Figure 1), meaning that they were genetically more distant. These less thermotolerant isolates also had different microsatellites patterns. The observed differences in thermotolerance and thermal growth curve shape might be explained by the different environmental and elaboration conditions of pulque and henequen mezcal. The Mexican central plateau where pulque is produced has a warm-temperate, subtropical climate with mild winters while Yucatan has a tropical climate with high temperatures throughout the year. Moreover, contrary to pulque, henequen mezcal elaboration involves high temperatures. Although further studies are required to understanding the physiological basis of these differences in thermotolerance, it can be speculated that *K. marxianus* populations derived from henequen mezcal elaboration process evolved to adapt to higher temperature niches and other harsh conditions. More studies are needed to understand the bases of these differences.

Tolerance of the isolates to CFW at different temperatures was tested to infer the relationship between thermotolerance and cell wall structure. Here, Kmx11 and Kmx15 from pulque were highly resistant to CFW at 30 and 42 °C (Figure 5) indicating that they had a lower chitin content maybe explaining their lowest thermotolerance. On the contrary, the isolates from henequen presented a lower CFW tolerance indicating they had more chitin in their cell walls. The fact that the CFW tolerance of these isolates decreased at 42 °C may indicate that chitin content in the cell wall was higher at 42 °C or that chitin was more accessible. Kmx22 from fermented cooked henequen juice presented the lowest tolerance to CFW indicating it had the highest chitin content among all isolates or that the chitin was more exposed to the dye. The results obtained somehow indicate that there is a relationship between the cell wall structure and thermotolerance in *K. marxianus* although the involvement of cell wall in the adaptation and tolerance of yeasts to temperature has received a lot of attention [81]. Detailed cell wall structural studies are needed to confirm the role of chitin and its cross-linking with other polysaccharides in the thermotolerance of *K. marxianus*.

The isolates were also evaluated against stress conditions relevant for the use of lignocellulosic hydrolysates as carbon sources to produce bioethanol and other products (Table 4) [54,82,83]. Osmotolerance was evaluated using glucose as the solute. All the isolates, except Kmx15 from pulque whose tolerance limit was lower at 42°C, tolerated a maximum of 30% of glucose. This tolerance limit is low compared to *S. cerevisiae* and other yeast species that can at least tolerate 50% glucose at 30°C [82]. Concerning ethanol tolerance, the isolates from pulque and henequen mezcal tolerated between 5 and 10% ethanol at 30°C which is similar to other non-conventional yeasts, but low in comparison to *S. cerevisiae*, the most ethanol tolerant yeast species (14% of ethanol) [82]. Contrary to osmotolerance, ethanol tolerance decreased at 42°C in all isolates, confirming that temperature and ethanol exert a synergistic toxic effect on yeast cells, mainly by disrupting the plasma membrane integrity [84]. As previously reported, the isolates were more tolerant to K^+^ than to Na^+^ [54,82]. The two isolates from pulque, Kmx11 and Kmx15, were completely intolerant to NaCl at 42°C. It can be assumed that this effect was mainly due to the Na^+^ ion toxicity rather than to general osmotic stress as these isolates were tolerant to 30% glucose at 42°C. As shown by CFW susceptibility assays, Kmx11 and Kmx15 had a different cell wall structure which may limit their mechanical resistance to ionic osmotic stress. With regard to fermentation inhibitors, it has been reported that acetic acid tolerance is variable among *K. marxianus* and that this yeast, as *S. cerevisiae*, is more sensitive to acetic acid as temperature increases [56,81]. Acetic acid tolerance of the isolates reported here was low (1.5 g/L at at 42°C) compared to the most tolerant strain reported to date, *K. marxianus* CECT 10875, which can tolerate up to 10 g/L acetic acid at 42°C. Acetic acid tolerance of *S. cerevisiae* is also highly variable, from 0.6 to 12 g/L, depending on the pH and composition of the culture medium [85]. All the isolates of this study displayed a good furfural tolerance, between 1. 5 to 2 g/L, considering that the most furfural tolerant *S. cerevisiae* strains found in a collection of 71 environmental and industrial isolates were able to grow in the presence of up to 3 g/L furfural [86]. High temperature did not affect furfural tolerance on a negative way, on the contrary, tolerance to furfural was maintained or even increased at 42°C. This effect may not be surprising considering that furfural detoxification to the less toxic furfuryl alcohol is NADPH-dependent [87] and that *K. marxianus* produces more NADPH at high temperature to fuel antioxidant systems that scavenge the reactive oxygen species formed under heat stress [88]. A similar effect was observed for the phenolic fermentation inhibitor CA except for the two pulque isolates Kmx11 and Kmx15, which tolerated less CA at 42°C than at 30°C. CA has been identified as the most toxic phenolic compound derived from lignin in lignocellulosic hydrolysates [83]. Albeit their significant toxicity at low concentrations, phenolic compounds are the less studied group of inhibitors [89]. The mechanisms of yeast tolerance to phenolic compounds are *in situ* reduction to less toxic alcohols, cell wall remodeling and protein homeostasis [89]. It has also been shown that CA can act as a cell wall active agent with antifungal activity [90]. Differences in the cell wall structure of the pulque isolates may explain their lower tolerance to CA.

Some biomass feedstocks also contain heavy metals as zinc, cadmium, manganese and copper [83] which are essential elements in small quantities but toxic for cells in excessive amounts. The tolerance of the isolates was comparable to that of *S. cerevisiae* in the case of zinc and cadmium [54] but higher in the case of copper. Considerable intra and interspecific variations in Cd and Cu tolerance has been found among 15 yeast species (not *K. marxianus*) from water, soil and plant environments [91]. Little is known so far about the different mechanisms underlying metal tolerance in yeasts. In the case of Cd, the pulque isolates and strain Kmx22 were the least tolerant to this heavy metal. These isolates also presented different behaviors in the presence of the cell wall perturbing agent CFW so, differences in cell wall structure may be related to their lowest Cd tolerance.

The *K. marxianus* isolates were finally evaluated for their ability to produce ethanol from the cellulosic fraction of a corncob hydrolysate by SSF. The obtained ethanol yields (Table 5) were similar to those already described for *K. marxianus* CECT 10875 in the SSF of different lignocellulosic biomass hydrolysates in shake flasks fermentations as in the present work [11]. The most performant isolate was Kmx11 with an ethanol yield of 56.4% and a residual glucose concentration of 0.8 g/L, followed by Kmx16, Kmx21, Kmx22, Kmx24, Kmx14 and Kmx15 which showed the lowest ethanol yield (43.3%) and the highest residual glucose (15 g/L) at the end of the fermentation trials (Table 5). Acetic acid concentration was almost constant during the SSF and corresponded to the concentration initially present in the hydrolysate (around 3 g/L). Although the isolates could not tolerate 3 g/L of acetic acid in assays performed on YPD agar at 42°C, they could efficiently ferment the corncob hydrolysate. It has been reported that results obtained from assays in laboratory media and synthetic hydrolysates generally cannot be extrapolated to the fermentation of real lignocellulosic hydrolysates due to the complexity of their composition, the presence of solids in high concentrations, minerals, and other nutrients or antinutrients [92]. It has been reported that, despite the inhibitory conditions present in wheat straw and sugarcane bagasse hydrolysates, a *K. marxianus* strain (SLP1) isolated from an *Agave salmiana* mezcal must showed a better performance than the *S. cerevisiae* Ethanol Red strain, currently utilized in the second-generation ethanol plants to produce ethanol from these hydrolysates [93]. Therefore, the isolates described here could be used as cell factories for the valorization of lignocellulosic biomass.

## 5. Conclusions

This study provides a general survey of the genetic and physiological diversity of *K. marxianus* isolates obtained from the elaboration process of pulque and henequen mezcal, two agave-derived alcoholic beverages. Significant genotypic and phenotypic diversity were found between the pulque and henequen mezcal isolates and among the henequen mezcal isolates, suggesting that local selective pressure may originate different *K. marxianus* populations in these environments. The differences in thermotolerance, cell wall structure, sugar assimilation profiles, and stress tolerance between pulque and henequen mezcal isolates could be related to differences in the climatic and process conditions as well as available carbon sources in the sampled substrates. The ability to ferment lactose and inulin and assimilate xylose, lactose, inulin, and cellobiose and produce ethanol at high temperatures represent interesting features for industrial applications. Further genomic and physiological studies are required to expand our knowledge on the intraspecific diversity and physiology of *K. marxianus* incorporating more non-dairy specimens and expanding the industrial use of this interesting species.

## Supplementary Materials

Tables S1 and S2.

## Author Contributions

“Conceptualization, PLO, MA, GB and SLB; methodology, PLO, MA, SDSR, AKCP, LP, GB and SLB; software, MA, SDSR and GB; validation, MA, GB and SLB; formal analysis, PLO, MA, GB and SLB; investigation, PLO, MA, SDSR, AKCP, LP, GB, SLB; resources, PLO and SLB; data curation, PLO, MA, GB and SLB; writing—original draft preparation, PLO, MA, GB and SLB; writing—review and editing, PLO and SLB; supervision, SLB; project administration, SLB; funding acquisition, PLO, LP and SLB. All authors have read and agreed to the published version of the manuscript.”

## Disclaimer/Publisher’s Note

The statements, opinions and data contained in all publications are solely those of the individual author(s) and contributor(s) and not of MDPI and/or the editor(s). MDPI and/or the editor(s) disclaim responsibility for any injury to people or property resulting from any ideas, methods, instructions or products referred to in the content.

