## Supplemental Table 1 for "Genotypic and Phenotypic Diversity of *Kluyveromyces marxianus* Isolates Obtained from the Elaboration Process of Two Traditional Mexican Alcoholic Beverages Derived from Agave: Pulque and Henequen (*Agave fourcroydes*) Mezcal"

Table S1. Carbon and nitrogen compounds assimilation patterns of the *K. marxianus* isolates obtained from the elaboration process of henequen mezcal and pulque. The two last columns show the results reported for *K. marxianus* in the literature for comparison.

| **Test** | **Isolates** | | | | | | | **Barnett et al. (2000)** | **Lachance (2011)** |
|---|---|---|---|---|---|---|---|---|---|
| | **Kmx11** | **Kmx14** | **Kmx15** | **Kmx16** | **Kmx21** | **Kmx22** | **Kmx24** | | |
| **Assimilation of carbon compounds** | | | | | | | | | |
| D-Galactose | + | + | + | + | + | + | + | + | + |
| L-Sorbose | - | - | - | - | - | - | - | +,- | - |
| D-Glusosamine | - | - | - | - | - | - | - | - | - |
| L Arabinose | - | + | - | + | + | + | + | +,- | v |
| D-Arabinose | - | - | - | + | - | - | - | - | - |
| Methyl α-D-glucoside | - | - | - | - | - | - | - | +,- | - |
| Salicine | + | + | + | + | + | w | + | +,- | v |
| Arbutin | + | - | + | + | + | + | + | +,- | nd |
| Melibiose | - | - | - | - | - | - | - | - | - |
| Melezitose | - | - | - | - | - | - | - | +,- | - |
| Starch | nd | - | nd | - | - | - | - | +,- | - |
| Erythritol | - | - | - | - | - | - | - | - | - |
| D-Glucitol | D | + | + | + | d | w | + | +,- | v |
| Galactitol | - | - | - | - | - | - | - | - | - |
| myo-Inositol | - | - | - | - | - | - | - | - | - |
| 2 Keto D gluconate | - | - | - | - | - | - | - | - | - |
| D-Gluconate | - | - | - | - | - | - | - | - | - |
| D-Glucuronate | - | - | - | - | - | - | - | - | - |
| Methanol | - | - | - | - | - | - | - | - | - |
| **Assimilation of nitrogen compounds** | | | | | | | | | |
| Nitrate | - | - | - | - | - | - | - | - | - |
| Nitrite | - | - | - | - | - | - | - | - | - |
| Ethylamine | + | + | + | + | + | + | + | + | + |

| | | | | | | | | | |
|---|---|---|---|---|---|---|---|---|---|
| L-Lysine | **+** | **+** | **+** | **+** | **+** | **+** | **+** | **+** | **+** |
| Cadaverine | **-** | **+** | **+** | **+** | **+** | **+** | **+** | **+** | **+** |
| Creatine | **-** | **-** | **-** | **-** | **-** | **-** | **-** | **-** | **nd** |
| Creatinine | **-** | **-** | **-** | **-** | **-** | **-** | **-** | **-** | **nd** |
| **Thermotolerance** | | | | | | | | | |
| 37°C | **+** | **+** | **+** | **+** | **+** | **+** | **+** | **+** | **+** |
| 40°C | **+** | **+** | **+** | **+** | **+** | **+** | **+** | **+,-** | **+** |
| 42°C | **+** | **+** | **+** | **+** | **+** | **+** | **+** | **+,-** | **nd** |
| 45°C | **+** | **+** | **+** | **+** | **+** | **+** | **+** | **+,-** | **nd** |
| 48°C | **-** | **+** | **-** | **+** | **+** | **+** | **+** | **nd** | **nd** |
| 50°C | | **+** | | **+** | **+** | **-** | **-** | **nd** | **nd** |
| 52°C | | **-** | | **+** | **-** | | | **nd** | **nd** |
| 55°C | | | | **w** | | | | **nd** | **nd** |
| 56°C | | | | **-** | | | | **nd** | **nd** |
| **Tolerance to ethanol Checar art Sylvie** |  |  |  |  |  |  |  |  |  |
| 5% | **+** | **+** | **+** | **+** | **+** | **+** | **nd** | **nd** | **nd** |
| 6% | **+ M** | **+** | **+M** | **+** | **+** | **+** | **nd** | **nd** | **nd** |
| 7% | **+** | **+** | **+** | **+** | **+** | **+** | **nd** | **nd** | **nd** |
| 8% | **-** | **+** | **-** | **+** | **+** | **+** | **nd** | **nd** | **nd** |
| 9% | | **+** | | **+** | **+** | **+** | **nd** | **nd** | **nd** |
| 10% | | **-** | | **-** | **-** | **-** | **nd** | **nd** | **nd** |

+ positive; **-** negative; **w** weak, d delayed, **v** variable: positive response; **nd** non determined
