## Supplemental Table 2 for "Genotypic and Phenotypic Diversity of *Kluyveromyces marxianus* Isolates Obtained from the Elaboration Process of Two Traditional Mexican Alcoholic Beverages Derived from Agave: Pulque and Henequen (*Agave fourcroydes*) Mezcal"

| | Kmx11 | | | | | | | Kmx14 | | | | | | |
|---|---|---|---|---|---|---|---|---|---|---|---|---|---|---|
| | **umax** | **stdv** | **λ** | **stdv** | **R2** | **Ymax** | **stdv** | **umax** | **stdv** | **λ** | **stdv** | **R2** | **Ymax** | **stdv** |
| **30 ºC** | 0.40 | 6.1% | 4.15 | 3.8% | 0.995 | 27.6 | 9.9% | 0.45 | 7.5% | 3.50 | 6.2% | 0.993 | 13.9 | 3.2% |
| **37 ºC** | 0.55 | 7.6% | 4.17 | 4.3% | 0.994 | 24.1 | 2.9% | 0.85 | 14.6% | 4.51 | 5.3% | 0.986 | 20.9 | 4.2% |
| **42 ºC** | 0.54 | 10.2% | 3.03 | 8.4% | 0.989 | 16.1 | 3.9% | 1.12 | 12.9% | 3.87 | 4.3% | 0.992 | 17.7 | 2.8% |
| **45 ºC** | | | | | | | | 1.11 | 16.1% | 3.54 | 5.8% | 0.987 | 17.5 | 3.4% |

| | Kmx15 | | | | | | | Kmx16 | | | | | | |
|---|---|---|---|---|---|---|---|---|---|---|---|---|---|---|
| | **umax** | **stdv** | **λ** | **stdv** | **R2** | **Ymax** | **stdv** | **umax** | **stdv** | **λ** | **stdv** | **R2** | **Ymax** | **stdv** |
| **30 ºC** | 0.63 | 12.2% | 4.28 | 6.0% | 0.986 | 18.7 | 4.7% | 0.49 | 6.8% | 3.41 | 5.40% | 0.995 | 13.4 | 2.8% |
| **37 ºC** | 0.68 | 6.9% | 4.39 | 3.1% | 0.996 | 20.0 | 2.3% | 0.51 | 10.0% | 4.16 | 6.08% | 0.990 | 23.6 | 4.0% |
| **42 ºC** | 0.58 | 10.5% | 2.81 | 6.1% | 0.997 | 11.8 | 3.6% | 0.91 | 7.7% | 3.85 | 3.13% | 0.996 | 17.3 | 2.0% |
| **45 ºC** | | | | | | | | 0.69 | 12.4% | 0.52 | 6.02% | 0.995 | 15.8 | 3.8% |

| | Kmx21 | | | | | | | Kmx22 | | | | | | |
|---|---|---|---|---|---|---|---|---|---|---|---|---|---|---|
| | **umax** | **stdv** | **λ** | **stdv** | **R2** | **Ymax** | **stdv** | **umax** | **stdv** | **λ** | **stdv** | **R2** | **Ymax** | **stdv** |
| **30 ºC** | 0.52 | 8.3% | 3.63 | 5.8% | 0.993 | 13.5 | 3.3% | 0.59 | 9.9% | 4.41 | 5.0% | 0.991 | 13.3 | 3.7% |
| **37 ºC** | 0.83 | 12.6% | 4.56 | 5.8% | 0.990 | 20.4 | 3.8% | 0.62 | 7.7% | 4.10 | 4.1% | 0.995 | 19.5 | 2.8% |
| **42 ºC** | 1.15 | 13.1% | 3.74 | 5.8% | 0.992 | 17.4 | 3.3% | 0.56 | 7.4% | 2.73 | 6.5% | 0.994 | 20.9 | 2.7% |
| **45 ºC** | 0.99 | 10.3% | 3.44 | 5.8% | 0.994 | 15.7 | 2.4% | 0.68 | 8.5% | 3.29 | 5.3% | 0.994 | 14.9 | 3.3% |

| | Kmx24 | | | | | | |
|---|---|---|---|---|---|---|---|
| | **umax** | **stdv** | **λ** | **stdv** | **R2** | **Ymax** | **stdv** |
| **30 ºC** | 0.43 | 4.1% | 3.40 | 3.7% | 0.998 | 11.6 | 1.8% |
| **37 ºC** | 0.84 | 9.9% | 4.57 | 3.0% | 0.997 | 20.6 | 2.4% |
| **42 ºC** | 0.80 | 13.0% | 3.84 | 5.9% | 0.989 | 15.7 | 3.8% |
| **45 ºC** | 0.93 | 10.5% | 3.86 | 4.2% | 0.994 | 14.4 | 2.7% |

| | | | |
|---|---|---|---|
| R2.avg | 0.993 | stdv.umax | 9.9% |
| R2.min | 0.986 | stdv.λ | 5.0% |
| R2.min | 0.998 | | |
